# Policy-Gradient Reinforcement Learning as a General Theory of Practice-Based Motor Skill Learning

**DOI:** 10.1101/2025.10.17.682587

**Authors:** Adrian M. Haith

**Affiliations:** Baltimore, MD, USA

**Keywords:** motor learning, motor skill, practice, reinforcement learning

## Abstract

Mastering any new skill requires extensive practice, but the computational principles underlying this learning are not clearly understood. Existing theories of motor learning can explain short-term adaptation to perturbations, but offer little insight into the processes that drive gradual skill improvement through practice. Here, we propose that practice-based motor skill learning can be understood as a form of reinforcement learning (RL), specifically, policy-gradient RL, a simple, model-free method that is widely used in robotics and other continuous control settings. Here, we show that models based on policy-gradient learning rules capture key properties of human skill learning across a diverse range of learning tasks that have previously lacked any computational theory. We suggest that policy-gradient RL can provide a general theoretical framework and foundation for understanding how humans hone skills through practice.

## Introduction

Learning a new motor skill invariably requires extensive, repetitive practice before we become proficient at it. Through practice, we gradually refine execution of motor skills through a combination of fine-tuning the exact motor commands we send to our muscles, and reducing the variability of these motor commands from movement to movement^1–3^. The learning principles underlying this gradual improvement are not clearly understood. Here, we show that practice-based motor learning across a variety of tasks is well captured by policy-gradient reinforcement learning, a simple and well-established model-free reinforcement learning (RL) method that is widely used in robotics and in many other continuous control settings^4–6^.

Most existing computational theories of motor learning are based on principles of *supervised learning*. According to these theories, observed task errors (e.g. missing a target to the left/right) are used to guide compensatory changes in motor output^7–9^. Doing this successfully, however, depends critically on pre-existing knowledge about the structure of the task in order to translate observed task errors into appropriate changes in motor commands^10–12^, but this knowledge may not always be available when learning a new skill. These *error-based* learning models account well for how people rapidly adapt already familiar well-practiced movements to compensate for imposed perturbations^9,13,14^, but cannot explain how brand new skills are learned and refined over hours of practice. Furthermore, error-based learning models account only for overall shifts in motor output and offer no account of the changes in variability that are a key characteristic of practice^2,15,16^. Alternative frameworks besides supervised learning are therefore needed to understand practice-based learning of motor skills.

Here, we show that motor skill learning through practice can be described using principles of *reinforcement learning* (RL), specifically, policy gradient methods. Policy gradient is a simple, model-free RL method that requires no prior task knowledge and guides improvements in performance based solely on observed states, actions, and a “reward” signal which provides a simple, scalar assessment of task success. We show how policy-gradient learning rules naturally account for changes in the mean and variance of behavior across a range of established skill-learning paradigms that have previously lacked any computational theory.

## Results

### Policy Gradient RL Framework

In general, RL describes the scenario in which an agent must learn from experience how to act in order to optimize an unknown reward function *r*. RL algorithms typically specify a *control policy* in the form of a probability distribution *π*_*θ*_(*a*|*s*), over potential actions *a*, given current task state *s*.

This policy depends on adjustable parameters *θ* (which typically describe the mean and variance of selected actions) and the goal of RL is to determine policy parameters *θ* that maximize the overall expected reward 𝔼[*r*(*s*, a)]. Pol*icy gradient* methods, attempt to optimize reward by following the gradient of the expected reward function with respect to the parameters 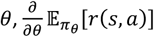. It is not possible to determine this gradient exactly, since the reward function is not known. However, the policy gradient theorem shows that this gradient equivalently be expressed as

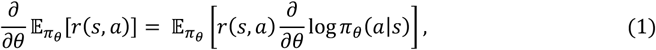

which, since _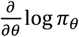_ is known, can be straightforwardly approximated in a Monte Carlo manner by sampling actions and associated rewards. If this gradient estimate is based on just a single sample of actions and outcomes from one trial, this leads to the following update rule:

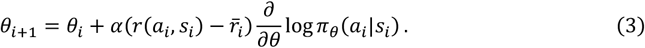

which describes how policy parameters *θ* on trial *i* should be updated based on the state *s*_*i*_, action *a*_*i*_, and reward outcome *r*(*a*_*i*_, s_*i*_). Note that here, the raw reward term *r*(*a*_*i*_, s_*i*_) in Equation (2) has been adjusted by 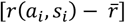, where 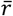is a running estimate of the recent rewards. This baseline subtraction is commonly employed in policy-gradient applications since it reduces the variance of the gradient estimate when using small numbers of trials.

This update rule, first introduced in 1992 under the moniker REINFORCE^5^, serves as the foundation of policy-gradient methods for RL. The exact form of the update depends on the nature of the policy distribution *π*_*θ*_(*a*_*i*_|*s*_*i*_) and its parameters *θ*. We consider policies that take the form of a multivariate Gaussian distribution over actions, with *π*(*a*|*s*) ∼ *N*(µ, Σ). In this case, the policy-gradient update for the mean µ becomes:

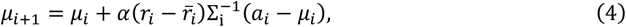

where µ_*i*_ and Σ_*i*_ are the policy mean and covariance on trial *i*.

This simple learning rule is illustrated in Figure 1A, which depicts an arbitrary reward function on a 2-D action space. The update rule in Eq. (4) shifis the mean of the distribution towards the sampled action *a*_*i*_, with a sign and amplitude that depends on the RPE, 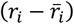. If the reward obtained is better than the average of recent previous attempts (i.e. if the RPE is positive), the policy mean µ will be updated towards action *a*_*i*_ (Fig. 1B). Conversely, if the outcome is worse than the average past behavior (negative RPE), the policy mean will become shifted away from action *a*_*i*_ (Fig. 1C).

**Figure 1.**
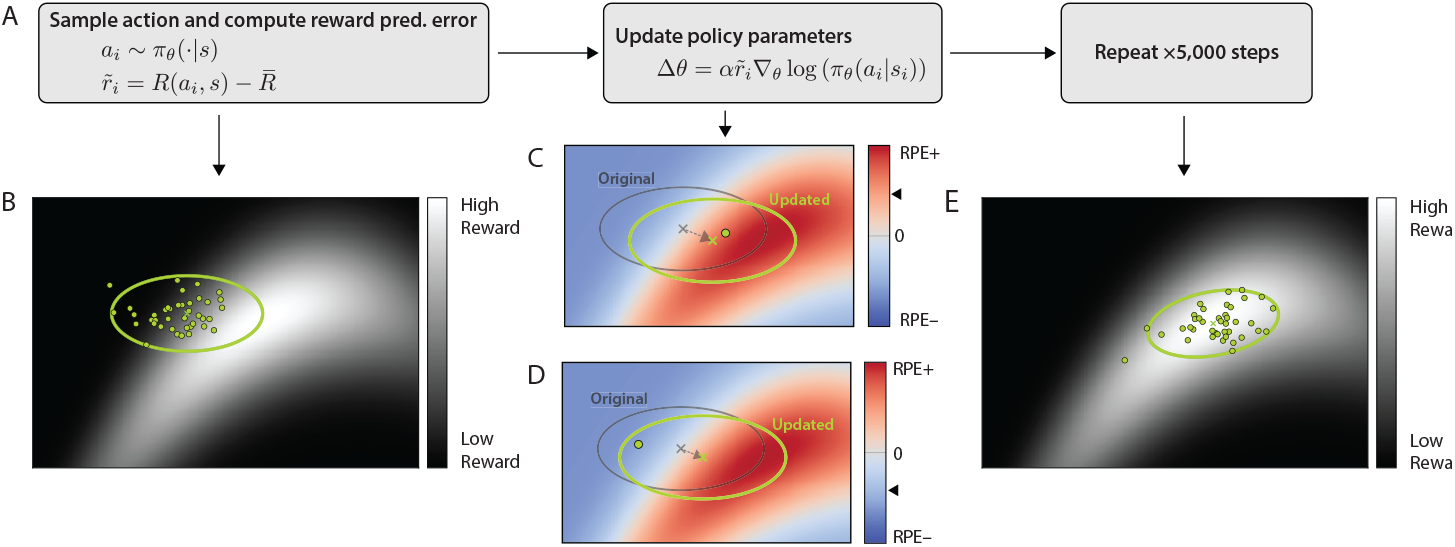
Illustration of policy-gradient RL. A) Outline of policy gradient algorithm. B) Reward landscape for a task with 2-dimensional action space, and actions generated according to an initial policy, in this case a 2-dimensional Gaussian. The green ellipse indicates the covariance of the Gaussian policy (90% confidence interval), while dots represent individual samples. C) Illustration of policy update for a positive reward prediction error. In this trial, the randomly selected action leads to a beKer-than-average reward (positive reward prediction error). Consequently, the mean is updated towards the sampled action. D) Policy update for a negative reward prediction error. Here, the randomly selected action results in a worse-than-average reward (negative reward prediction error). Consequently, the mean is updated away from the sampled action. E) After a few thousand samples, the policy converges on the region of greatest reward.

In addition to updating the mean of the policy, Eq. (2) can also be applied to derive a learning rule for the covariance matrix of the policy, Σ. This can be represented as a diagonal matrix of log-eigenvalues *v*:

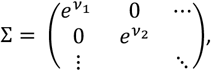

which ensures a valid covariance matrix for any value of (*v*_1_, *v*_2_). In this case, the general policy-gradient update for parameter *v*_*k*_ on timestep *i* becomes:

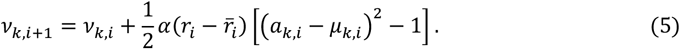

Parametrizing the covariance matrix in this way assumes that the actions along each dimension are sampled independently, but it is also possible to derived updates for learning covariance between actions.

Equation 3 above provides a general, trial-by-trial learning rule that can be applied to any task for which a parametrized, differentiable policy is specified. As a model of learning, its main free parameter is the learning rate *α* (though this can, in principle, take different values for different policy parameters). We show how this simple framework yields models of learning that align closely with human behavior across a range of very different tasks.

### Application to a skittles-toppling throwing task

We first applied the policy-gradient learning rules described above to simulate learning in the skittles-toppling task developed by Müller and Sternad^17^. This well-studied task is inspired by the real-world bar game of skittles in which participants must throw a ball, suspended by a string from a supporting central post, towards a target skittle, aiming to topple it (Fig. 2A). In laboratory versions of this task, it has been found that people gradually learn to improve their performance over hundreds to thousands of trials of practice^17,18^. Participants’ attempts on each trial are characterized by the launch angle and velocity of the ball, and only certain combinations of these actions will successfully topple the skittle (Figure 2B). Participants tend to initially attempt throws that follow a broad distribution in this action space but their actions become gradually refined with practice, first by shifting the mean of the distribution towards more successful actions, followed by more gradual reduction in the variability of actions and alignment with the shape of the reward landscape^2,17–19^ (Figure 1B).

**Figure 2.**
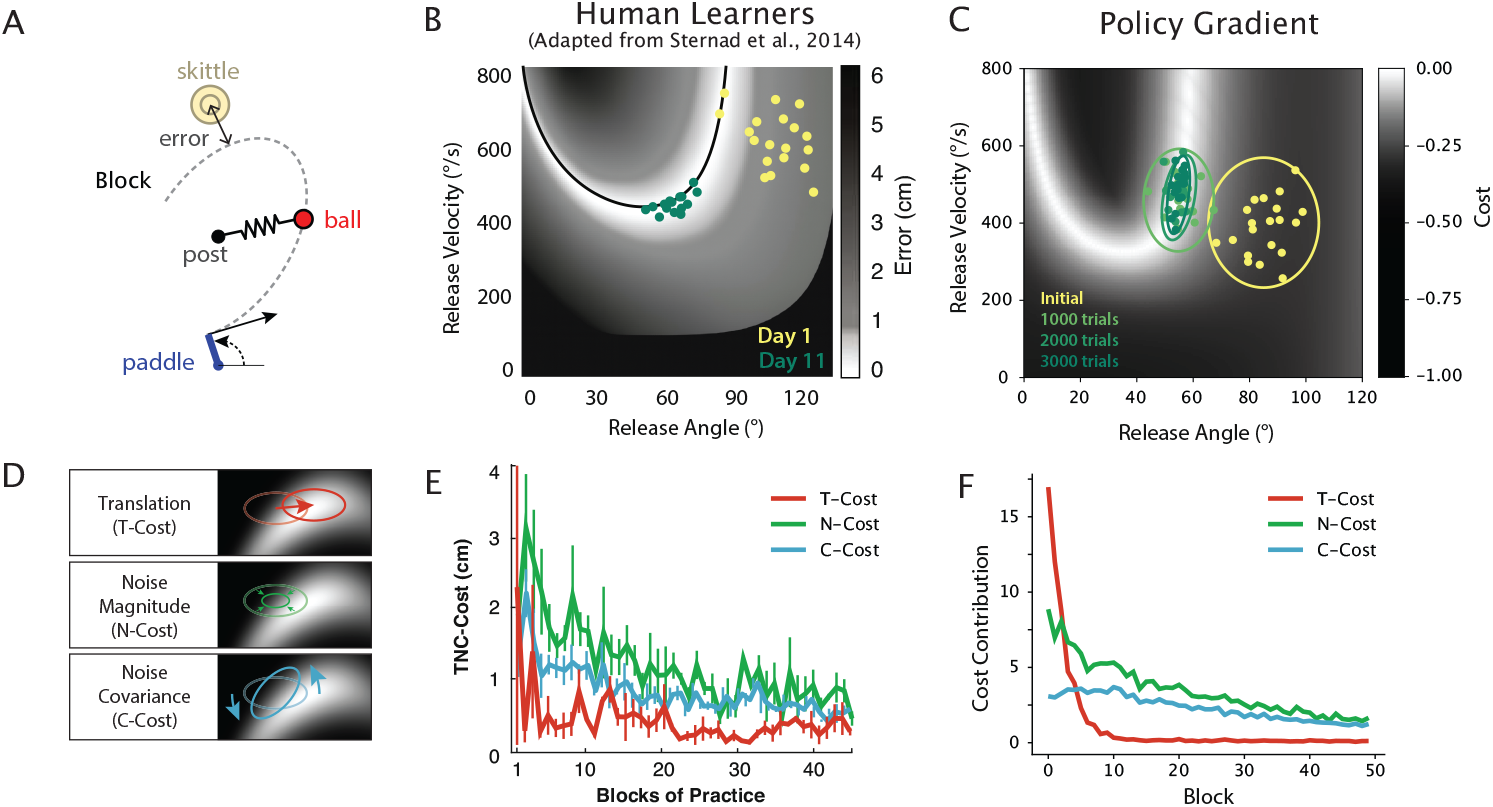
Policy-gradient RL model of a throwing task. A) In Sternad and colleagues’ virtual skiKles task^17,18^, participants swivel a paddle and press a buKon to release a ball, aiming to topple a skiKle while avoiding a central post. The ball’s movement is guided by an elastic force acting towards the central post, and is fully determined by the initial release angle and velocity. The task error is taken as the minimum distance of the balls trajectory to the center of the skiKle. B) Example human performance in this task: Heatmap shows task error given release angle and velocity, adapted from^18^. Yellow dots show example data from a human participant on their first day performing the task. Green dots show data from the same participant after 11 practice sessions (∼2,000 trials). C) Example performance of the policy gradient algorithm trained on the same task. Ellipses show 90% confidence regions for the policy at different stages of learning. D) Illustration of the TNC-Cost decomposition used to compare human and model behavior. Based on actions taken over a block of 60 trials, the *Tolerance Cost* (T-Cost) estimates potential performance gains achievable over the current policy by translating the policy in action space. The *Noise-Amplitude Cost* (N-Cost) estimates potential performance gains achievable by uniformly scaling the noise. The *Noise-Covariance Cost* (C-Cost) estimates potential performance gains achievable by optimizing the policy covariance. E) TNC-Cost decomposition^19^ for 9 participants, adapted from^18^. F) TNC-Cost decomposition for simulated learning of the policy-gradient RL model.

We modeled behavior in this task using the policy-gradient RL learning rule in Equation (2). As in the example in Figure 1, we represented the model as a multivariate Gaussian distribution with mean µ and covariance matrix Σ. In addition to representing the log-eigenvalues of the covariance matrix, we also introduced a further parameter that describes the covariation between actions through a rotation of the covariance matrix Σ. This resulted in five parameters in total: 2 for the mean and 3 for the covariance. We used a reward function equal to the minimum distance between the ball and the skittle, and used the same learning rate, *α* = 0.07, for all parameters (chosen so that simulations approximately matched the timescale of human learning). Similar to human participants, the policy-gradient RL model exhibited an initial shift in mean towards more successful actions, followed by a gradual reduction in variance and alignment of the covariance matrix with successful regions of the action space (Fig. 2C).

To compare behavior between our model and human learners in more detail, we applied a well-established analysis^18,19^ that characterizes the progression of different aspects of participants’ behavior. This analysis decomposes an individual’s performance in a given block into three distinct components by which it deviates from best-possible performance (Figure 2D): i) potential improvement obtained by translating the policy in action space (T-Cost), ii) potential improvement obtained by scaling the overall variance of the policy (N-Cost), and iii) potential improvement obtained by altering the covariance between actions (C-Cost). This analysis has revealed that human learning proceeds by rapid initial translation of the mean action (reductions in the T-Cost), accompanied by more gradual shrinking of variance (reductions in the N-Cost) and alignment with reward landscape (C-Cost), with a slightly greater contribution from the N-Cost (Figure 2E). The policy-gradient RL algorithm exhibited almost identical characteristics to human behavior (Figure 2F). The only major difference was that the model exhibited slightly slower reduction of the T-cost in the model compared to human learners. This indicates slower convergence on a suitable mean action and is likely attributable to a poorer initial policy, which was random for our model but in human learners could be informed by an initial intuitive understanding of the structure of the task. Nevertheless, the overall pattern of practice-based improvement was well-captured by policy-gradient RL.

In the skittles task, therefore, policy-gradient RL provided a very plausible and parsimonious model of how humans learn to improve their performance through practice, yielding successful learning over a similar timescale and with analogous characteristics to human learners.

### Application to a “de novo” cursor control task

We next applied the policy gradient learning rule to a “*de novo”* cursor control learning task. In this task, participants have to maneuver a cursor using movements of the left and right hand through a non-intuitive mapping (Figure 3A). This task requires participants to learn a new controller “*de novo*”, rather than by adapting an existing policy^20^. Participants initially find it extremely challenging to reach the targets with the cursor, but they gradually improve their performance over thousands of trials of practice across multiple sessions (Figure 3C).

**Figure 3.**
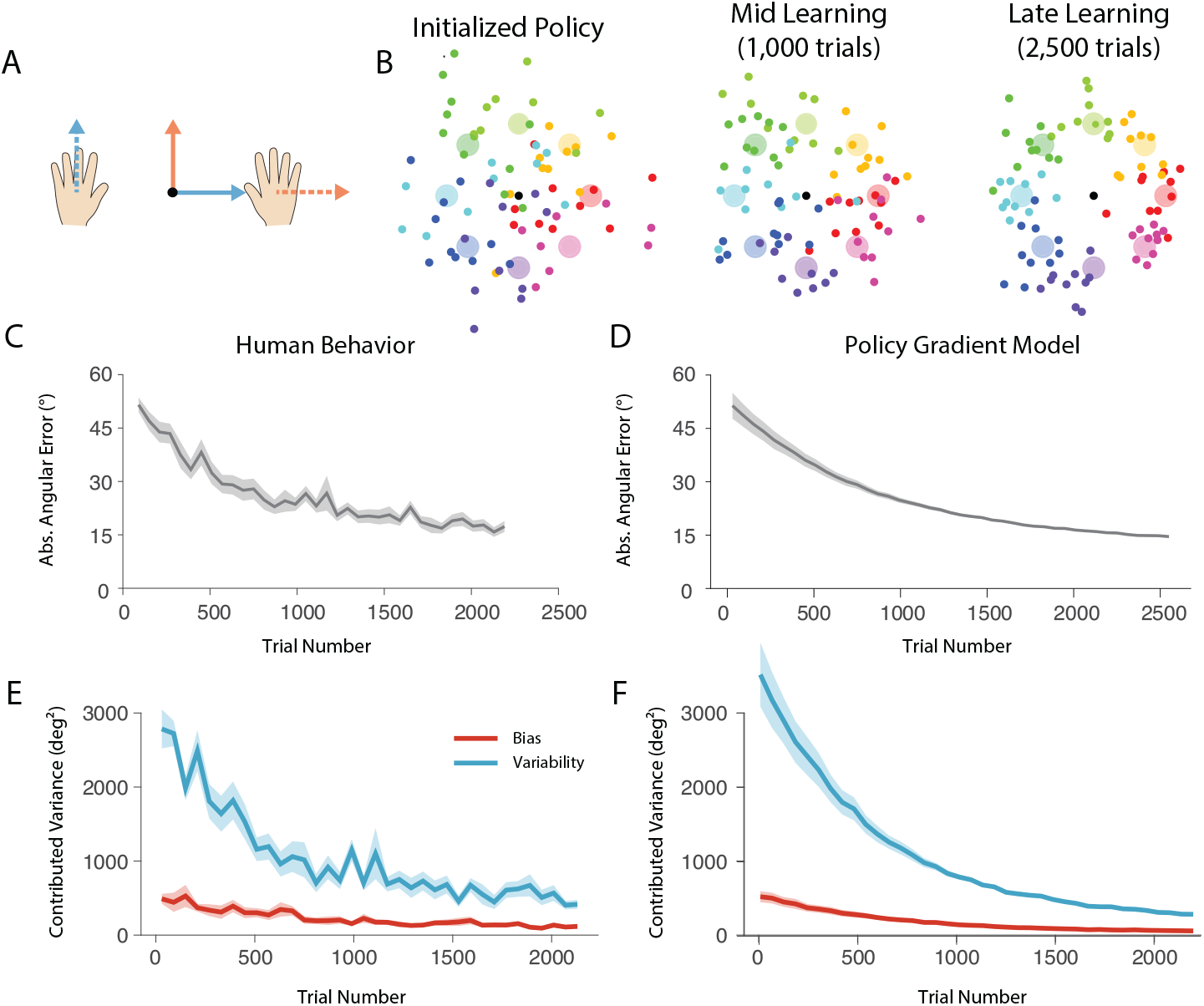
Policy-Gradient RL model of learning a cursor-control task. A) In the bimanual cursor control task, participants must learn to control a cursor using a non-intuitive mapping from the positions of their left and right hands to an on-screen cursor. Forward-backward movement of the left hand leads to left-right movement of the cursor, and left-right movement of the right hand leads to forward-backward movement of the cursor. B) Learning of a policy-gradient reinforcement learning model for this task. Large circles represent different target locations, to be reached by the cursor from a central starting location. Each dot represents a trial endpoint, with color indicating the associated target for that trial. The three panels illustrate the policy at the outset of training, after 1,000 training trials, and after 2,500 training trials. C) Human performance in this task. Each curve shows average absolute directional error across blocks of 60 trials, averaged across N=13 participants. Shaded region indicates +/-standard error in the mean across participants. D) Average performance of the policy-gradient RL models with policies initialized to match initial behavior of human participants. E) Bias-variance decomposition for human performance over trials, averaged across participants. Most practice-based improvement is aKributable to reductions in variability. F) Averaged bias-variance decomposition for policy-gradient RL models with policies initialized to match initial behavior of human participants. Shaded regions indicate +/- sem across participants (E) or across models initialized to different participants (F).

To model behavior in this task using policy-gradient RL, we assumed that individuals generated point-to-point movements of their left and right hands, and systematically varied the direction and extent depending of these movements depending on the target location. The action space was therefore 4-dimensional. To allow for a policy where the actions could vary based on the target angle (state, *s* ), the mean action was constructed as a weighted sum of local basis functions, µ(*s*) = ?_-_ *w*_-_*Φ*_-_(*s*), yielding a mean action that depended smoothly on the state. We updated the weights *w*_-_ based on the learning rule derived from Equation (4). We allowed the estimate of recent reward, 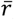, driving the learning to also depend on the target direction (state *s*) using the sam e basis functions (making it a *value function* in RL terminology). In addition to learning suitable mean actions, we assumed distinct (state-independent) variances σ_*k*_ along each dim ension of the action space, and also learned these parameters via policy-gradient updates according to Equation (5). As a cost function, we used the distance between the end cursor location and the center of the target, and w e set the learning rate to *α* = 0.1 in order to approximately match the timescale of human behavior.

Despite the added complexity of the policy being target-dependent and in a higher-dimensional, redundant action space, the policy-gradient model was able to learn the task within the same tim escale as human participants – around 2,500 trials^20^ (Fig. 2B). Initial performance of the policy-gradient model, which was initialized with random weights, was poor, however. Performance, assessed in terms of absolute initial directional error, did not reach initial hu man performance levels until after around 1,000 trials of practice. After this point, however, learning of the model resembled human learning quite closely. Therefore, although initial performance of human learners was far better than the random initializations used in our policy-gradient RL model, likely due to people’s explicit understanding of the structure of the task, the gradual, practice-based improvement over the next 2,000 trials may have occurred through policy-gradient RL.

To overcome the discrepancy in initial performance between the policy-gradient model and human learners, and thereby better compare the nature and time course of practice-based improvement, w e set the policy-gradient RL model’s initial policy to match early policies of individual hu man participants, estimated based on their behavior in the first 120 trials. We then simulated learning via policy-gradient given that subject-specific initialization. The learning simulated this way was closely aligned with human behavior, seen in the averaged time course over which the initial directional error improved with practice (Figure 3C,D).

We further analyzed human behavior in more detail by decomposing the total variance of participant’s initial angular errors into a component due to bias of their movements (squared average angular deviation from the target) and a component due to variability around the mean deviation. Fig. 3E shows the contribution of these two components, revealing that the bulk of participants’ improvement occurred through reduction of the variability of their movements. We applied the same analysis to learning simulated under the policy-gradient RL model and found that, with suitable parameter settings (*α*_μ_ = 0.1, *α*_)_ =0.02), the time course and nature of improvement were closely aligned with human behavior (Fig. 3F).

Therefore, similar to the skittles task, the practice-based component of learning in this task was well-captured by the principles of policy-gradient reinforcement learning.

### Application to a precision motor execution task

In the first two tasks we considered, the learning challenge is clearly one of action *selection*, and these tasks are thus are naturally modeled within the RL framework which is fundamentally concerned with learning to *select* suitable actions. In contrast, many other practice-based motor learning tasks are thought to require learning to *execute* actions better, a capability referred to as “motor acuity”, by analogy to visual acuity. One example of such a task is the “arc task”^16^, in which participants use wrist movements to maneuver an on-screen cursor through an intuitive but unpracticed mapping, similar to using a laser pointer. Participants must guide the cursor through a narrow, semi-circular channel to reach a target circle (Fig. 4A). Initially, participants’ trajectories are very variable, with a high proportion of failed trials in which the cursor leaves the channel. After around 1,000 trials of practice over multiple days, however, participants gradually become able to reduce the variability of their actions to improve the proportion of trials that remain inside the channel^16^ (Figure 4A, right panel).

**Figure 4.**
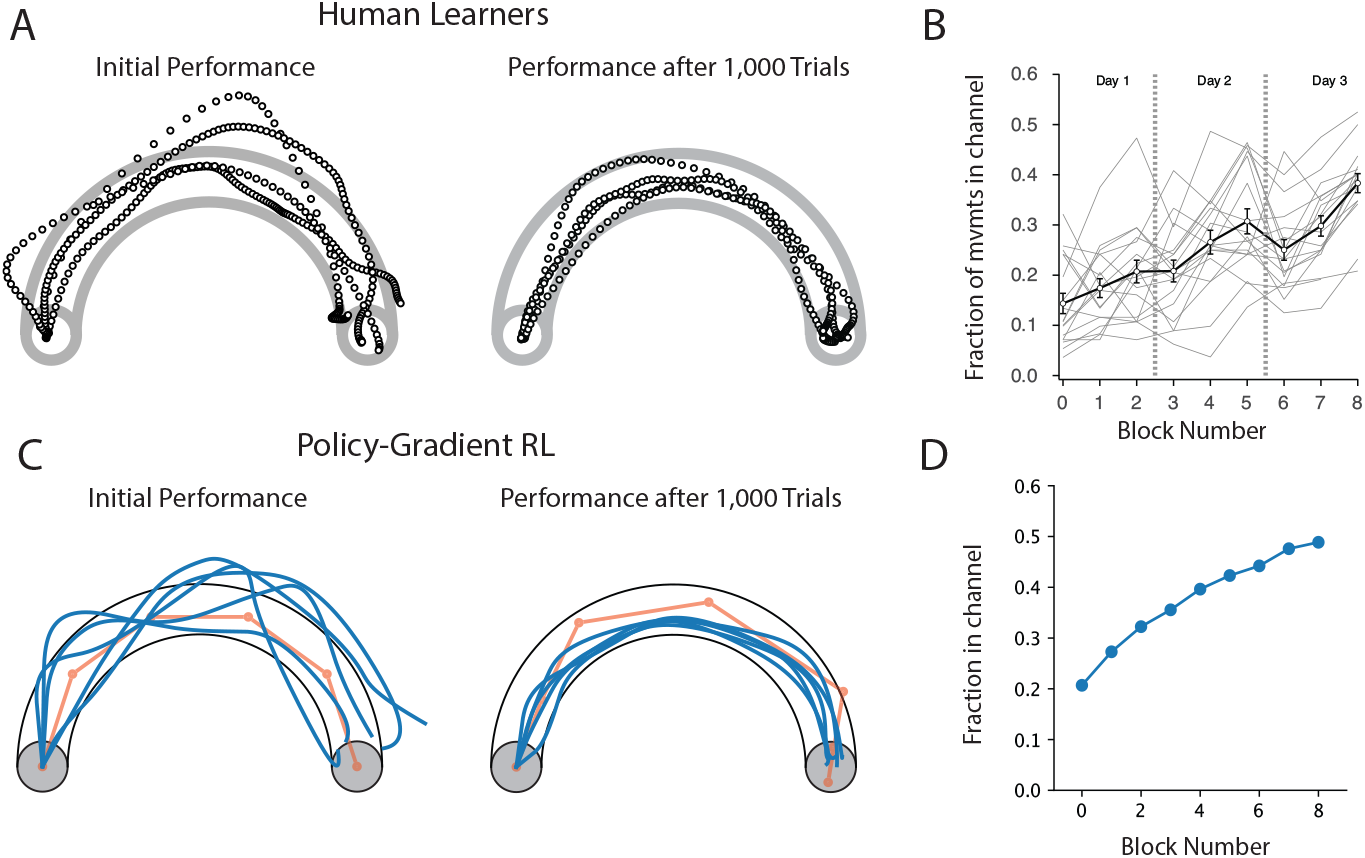
Policy-gradient model of learning to generate precise movements. A) Precision execution task, adapted from^16^. Participants used movements of their wrist to rapidly guide a cursor through an arc-shaped path from left to right with the goal of not leaving the channel. Initially, participants’ paths were highly variable and most trials were unsuccessful. After 8 sessions of practice (∼1,000 trials), participants’ movement variability was reduced, and the rate of successful movements increased. B) Fraction of trials successfully remaining within the channel throughout across 8 blocks of practice (120 trials per block), averaged across human participants. Panels A) and B), reproduced from^16^. C) Policy-gradient RL model of motor acuity learning. Initial policy was set to approximately match initial human performance. After 1,000 training trials, performance of the model was substantially improved. Red dots indicate intermediate goal locations used to model generation of the movement trajectories. The parameters of the policy were the 2-D location of the 6 goal locations, along with variance of their locations across trials. D) Fraction of trials staying within the channel, averaged across 100 simulated experiments.

Because the average cursor path does not change much and only the variability across attempts changes substantially with practice, this task has been thought to purely assess how people learn to *execute* a movement better with practice, reducing their motor execution noise. However, since the timescale of learning (around 1,000 trials) is not too different from that in the skittles and cursor control tasks, we wondered whether learning in this task might in fact be better understood as one of action selection. Although the required path for the cursor appears obvious, the mapping between wrist movements and the cursor location was not very familiar to participants and they could not have known the exact movements required. We therefore developed a model of learning in this task to test whether practice-based improvements in performance could be accounted for by the same learning principles that could explain practice-based improvements in the skittles and cursor-control tasks.

To simulate execution of individual movement trajectories in this task, we adopted a model in which movements are generated based on a pre-planned sequence of subgoals which are updated every 130 ms. This previously proposed control scheme is consistent with observed micro-structure of behavior when generating complex trajectories^16,21^. Therefore, although the movements themselves are continuous, they can be conceived as being generated based on a discrete sequence of 2-D subgoals. We supposed that this sequence of subgoals may be the higher-level actions that must be selected and optimized in order to generate successful movements. Therefore, we developed a policy-gradient RL model in which the action *a* specified the pre-planned sequence of subgoals (six 2-d goal locations for a movement lasting 780 ms), giving a 12-dimensional action space. The policy took the form of a Gaussian distribution over this action space, with 12 parameters specifying the mean of the control points (six 2-D locations in space), and a further 12 parameters specifying their variance along each axis (log-eigenvalues of a diagonal covariance matrix). We used a reward function that combined a number of considerations related to task success: reaching the correct endpoint, remaining inside the channel, and maintaining a smooth trajectory. We used a fixed learning rate of *α* = 0.2 for both mean and variance parameters, selected to approximately match the time course of human behavior.

Trajectories generated by the model looked qualitatively similar to those exhibited by human participants performing this task (Fig. 4C). Over 1,000 trials, the trajectories became progressively less variable until they most remained within the channel. We compared the time course of improvement over 8 blocks of 120 trials each between human participants (data from^16^), and our policy-gradient RL model, with the learning rate set to *α* = 0.2. The model paralleled participants’ steady improvement over the course of 8 blocks (960 trials).

Therefore, the policy-gradient RL framework not only successfully describes learning in tasks that obviously represent an action-selection challenge, the same principles can explain learning in tasks that appear to stress action execution, suggesting that apparent improvements in execution may in fact really be improvements in action selection driven by policy-gradient RL. This suggests that policy-gradient RL may be a very general principle underlying practice-based improvements in skilled motor performance.

## Discussion

Despite many decades of effort developing computational models of human motor learning, we have no clear theory of how most everyday skills are learned. Here, we have shown that a single learning rule (Equation 2) based on policy-gradient RL accounts well for patterns of practice-based improvement in a diverse range of motor skill learning paradigms. We therefore suggest that policy-gradient RL provides a plausible and promising general theory of practice-based motor skill learning.

Reinforcement learning has long been thought to be important for motor learning^22^, but few concrete computational models have been proposed. In previous work, RL-inspired algorithms have been applied to explain human motor learning in highly simplified tasks. Several studies have examined learning in 1-dimensional learning tasks in which subjects must learn to generate a specific action (e.g. reaching in a specific, but unknown, direction) given success/failure feedback about their performance^23–27^, and computational models have been proposed to account for behavior in these tasks. These models can mostly be construed as variants or approximations of policy-gradient learning rules^24–26^. Policy-gradient-like learning rules have also been successfully applied to explain locomotor adaptation^28^.

An important difference between the models presented here and previous work on reinforcement-based motor learning is that previous experiments and theories have mostly emphasized binary success/failure feedback. Learning to adapt one’s behavior under binary feedback is extremely challenging^29^, and becomes particularly difficult in higher-dimensional tasks^30^. In the models presented here, learning is driven instead by a scalar reward function that takes continuous values reflecting the relative quality of movement. Importantly, this scalar reward function might not directly reflect overt rewards associated with success or failure, like whether the skittle was toppled or not, but reflect a more continuous grading of outcomes, like distance to the skittle. While overt rewards have been found to influence the rate of motor learning in certain tasks^31–33^, these overt rewards appears to exert more of a modulatory effect, rather than serving as the main signal that drives learning.

One consistent finding in reward-driven motor learning tasks is that participants actively modulate their variability, increasing variability following errors^24,34,35^, seemingly to promote exploration. This increase in variability is not predicted by the basic policy-gradient RL update in Equation 2. However, it is possible that this phenomenon is an artifact of using tasks in which learners must rely only on binary feedback. With binary rewards, outcomes from a single trial carry little information about the gradient of the overall reward function and increased variance can improve a learner’s ability to reliably estimate the gradient of the reward function given few actions. With continuous rewards, there is less benefit to increasing variability, since even small variability in actions can carry reliable information about the direction of the reward gradient, provided the relative differences in rewards for different actions can be reliably discriminated.

Motor learning is often considered to be a highly cognitive enterprise^36–39^, with learning thought to involve a combination of reasoning about possible motor solutions to a task, and refinement of promising solutions^40^. The policy-gradient RL models presented here offer a concrete theory of the refinement part of this process. In this theory, however, the refinement occurs through a very simple learning rule that appears to have little need for cognitive involvement. The importance of cognitive contributions in the tasks we considered is clearly apparent in early learning in both the skittles task and the bimanual cursor control task, where initial human performance far surpassed that of randomized initial policies used in our models. Policy-gradient RL models initialized with random policies could require hundreds of trials of training before catching up to initial human performance levels. After that point, however, subsequent learning of the model was closely aligned to human learning. The superior initial performance of human learners compared to the model was likely attributable to high-level understanding of the structure of the task and the ability to reason about potential solutions that would fare much better than picking an initial policy at random. The fact that the gap between human learners, and our simple policy-gradient RL models was only really apparent at the outset of learning suggests that cognitive contributions to motor skill learning may be restricted only to the very early phases of learning, and there may be little need for cognitive involvement for subsequent, practice-based improvement.

The limited role for cognition in practice-based learning is in contrast to many theories of motor learning, which posit that cognition serves to guide action selection throughout the course of learning^37^. It also appears to contradict extensive literature in skill acquisition which has argued that gaining expertise in complex skills requires *deliberate prac7ce* – a concerted and cognitively demanding approach to practice^41,42^. Can the notion of cognitively-demanding deliberate practice be reconciled with a policy-gradient RL theory of practice? One possible role for cognition in practice, within the framework of policy-gradient RL, is that it may help to shape the reward function. The exact nature of the reward function is a subjective aspect of our model, though in additional simulation (not shown), we found that basic properties of learning were preserved even if the exact form of the reward function was varied (e.g. using squared distance to the goal, rather than absolute distance). It is well established that primary rewards may not be the most efficient way to train an RL agent, and that alternative pseudo-reward functions constructed internally can be significantly more effective^43,44^. Intuitively, a ‘good’ outcome of a golf swing at the driving range would look very different for a novice versus an accomplished golfer. The reward function therefore ought to be adjusted throughout the course of learning in order to maximize learning efficiency, in contrast to the static reward functions used in our models. This progressive reshaping of the reward function could be achieved through explicit, cognitive processes that oversee learning and assume the role of ‘critic’ (in the sense of actor-critic RL), providing the reward signal driving learning in Equation 2. This ‘cognitive critic’ proposal is consistent with the emphasis in deliberate practice on ensuring high-quality feedback on the quality of one’s performance and enhancing self-monitoring and assessment.

More broadly, the often slow nature of simple RL algorithms has led to the idea of meta-reinforcement learning^45,46^ – the problem of how to optimize learning speed of RL agents by constructing a suitable curriculum, potentially including adjustments to the reward function as well as selection of specific states and tasks to focus practice on. Cognition in skill learning, then, may more generally be required to accelerate the slow nature of RL, without actually being responsible for reasoning about action selection other than at the very outset. These influences of cognition may also account for the often substantial individual differences in learning across individuals in skill learning, rather than differences in learning rate parameters.

Having a concrete, computational theory of motor skill learning will be beneficial for gaining greater clarity in identifying its neural basis. The policy-gradient model in general requires two distinct sets of computations, one to compute the reward prediction error (or, more generally, the value function or advantage function) and one to update the policy. It is well known that dopaminergic neurons seem to compute a reward-prediction error, and are thus a clear candidate for computing the advantage function. Reward-related signals are observed throughout the brain^47^. Plasticity in the cortex is well known to be modulated by dopamine and dopaminergic projections to motor cortex are necessary for learning dexterous skills in rats^48–50^. Therefore, the policy itself is likely represented in motor cortical regions.

In conclusion, our results establish a concrete computational model of practice-based skill acquisition in humans that can describe learning across a diverse range of learning tasks that emphasize the importance of repeated practice. Given that policy-gradient methods are at the heart of recent advances in robotics^4,51^, and have also been successfully applied to musculoskeletal control problems^52^, it seems promising that policy-gradient RL could prove to be a universal principle that can account for the seemingly limitless range of skills that humans are capable of learning.

## Methods

### General Approach

For simplicity, we consider only single-timestep tasks in which an action or actions are selected only once, at the start of the movement, and these actions directly give rise to a reward. (We do not consider temporally-extended tasks in which the later states depend on prior actions, though the general policy-gradient framework is very much extendable to that setting).

We assume that users select actions *a* stochastically, based on the current state *s*, according to a policy specified as a probability distribution over actions, *π*_*θ*_(*a*|*s*). Here, *θ* denotes parameters of the policy, which are adjusted through learning. For a given task, actions lead to an outcome which is then associated with a reward function *r*(*s*, a), providing a scalar metric of performance. The goal of reinforcement learning is to identify policy parameters *θ*^*^ that maximize the average reward.

Policy gradient methods aim to optimize performance by adjusting the parameters *θ* by gradient descent. That is, they seek to follow the gradient,

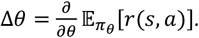

It is not immediately obvious how to calculate this expected gradient, since neither the expected reward nor its gradient with respect to *θ* is known. However the problem is simplified by the *policy gradient theorem*, which states that the gradient can equivalently be expressed as:

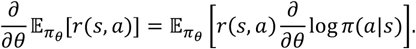

Critically, this expectation can now be approximated by sampling a large number of actions and associated costs, giving rise to a batch update rule as is typically used in reinforcement learning applications. It is possible, however, to use a batch size of 1, which yields an online learning rule which we adopt as our model of trial-by-trial skill learning. Therefore, on trial *i*, we assume that the policy parameters *θ* are updated according to

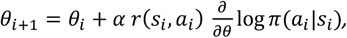

where *α* is a learning rate.

This learning rule can be adjusted to improve stability by replacing the reward in Eq.4.1 with a reward prediction error, i.e. subtracting out a baseline expected reward based on recent attempts:

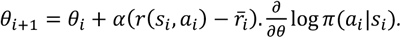

In our simulations, we update the average reward, _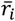_, trial-by-trial according to with β = 0.95.

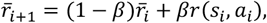

In all of our models, we assume that an individual’s policy *π*(*s*|*a*) follows a Gaussian distribution parametrized by a mean µ (which, may depend on the state *s*) and a covariance matrix Σ:

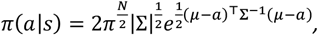

where *N* is the dimensionality of the action space. Updates to the mean in this case are given by:

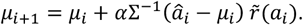

Policy-gradient methods are known to be vulnerable to instability triggered by occasionally large parameter updates. This issue is exacerbated when updates are based on single action-outcome observations, rather than batches, as is typical. A widely used method to avoid such instabilities is a method called proximal policy optimization^51^, which limits the size of parameter updates in a way that bound the possible ratio of probabilities of sampled actions between the original and updated policies. In line with this approach, we bounded updates to the mean to ensure that:

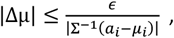

where ϵ is a small number, usually around 0.1. This corresponds to a linear approximation of the PPO bound for the mean of a Gaussian policy.

### Skittles task model

In the virtual skittles task, participants must launch a virtual ball towards a target skittle, while avoiding a central obstacle. The ball’s dynamics are governed by a springlike attractor towards the central obstacle. Participants’ actions in a given trial are the launch angle and velocity with which the ball is released, i.e., *a* = [*v, Φ*]^6^. The skittles task was modeled as described in^18^ and as illustrated in Fig. 2A. We assumed a paddle of length ***l***=0.4 m, situated 1.3 m below the central post, and with the skittle located 0.25 m to the right and 1.2 above the central post. We assumed a spring-like force attracting the ball towards the center of the task space, with stiffness ***k*** = **1** N/m, and with the ball’s mass equal to **0. 1** kg. To calculate the ball’s trajectory, we determined the initial release position and velocity of the ball:

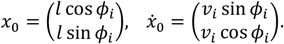

The full trajectory of the ball was then given by:

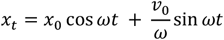

where 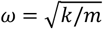. We determined the cost on a given trial by simulating the trajectory of the ball, and calculating the minimum distance between the ball and the center of the skittle, which served as the reward signal in our model:

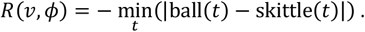

The learner’s policy was represented by a mean action µ = (*v, Φ*)^6^ and a covariance matrix Σ. To ensure that rotations of the covariance were meaningful despite the two different action dimensions being qualitatively different and potentially measured in different units, we represented the policy using a normalized, unitless co-ordinate system by subtracting the initial mean and dividing by the initial standard deviation for each action dimension:

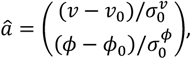

where *v*_8_ is the initial mean release velocity, *Φ*_8_ is the initial mean release angle, and 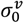 and 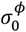: are the initial standard deviations. The initial normalized policy mean was then simply µ_8_ = (0,0)^6^ and with initial covariance matrix was given by the identity matrix, i.e. Σ_8_ = ***I***.

We parametrized Σ using an eigenvalue decomposition,

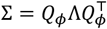

where *Q*: is a rotation matrix parametrized by angle *Φ*:

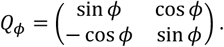

and Λ is a diagonal matrix of eigenvalues. To ensure numerical stability and avoid updates that could lead to negative eigenvalues (and an invalid covariance matrix), we represented this matrix in terms of its log-eigenvalues:

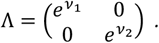

The overall policy was then specified by

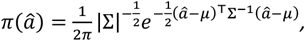

with five variable parameters in total, two for the mean Lµ_9_, µ_<_M, and three for the covariance matrix (*v*_1_, *v*_*_, *Φ*).

The policy gradient update is determined by the gradient of the log-probability. For the mean, this is given by

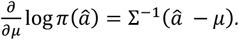

This leads to the following learning rule for updating the normalized mean action:

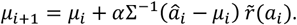

Here, _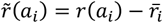_ is the reward prediction error, which depends on the average recent reward _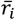_, which is updated after every trial according to:

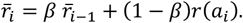

For the log-eigenvalues of the covariance matrix, the gradient of the log-probability is given by

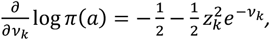

where 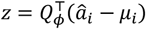 and *z*_+_ is its k^th^ element. This leads to the following learning rule for the log-eigenvalues *v*:

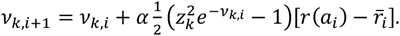

Lastly, for the orientation of the covariance matrix (given by the angle variable *Φ*), the gradient of the log-probability is given by

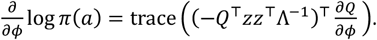

This leads to the learning rule:

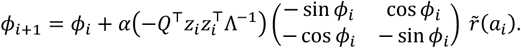

In all the above updates, *α* is a fixed learning rule shared across all parameters. Gradient calculations were confirmed using an automatic differentiation tool (pytorch).

For the TNC-Cost analysis, we grouped trials into blocks of 60 trials (to match human experimental data). For each block of 60 trials, we calculated the T-Cost, N-Cost and C-Cost as follows. For the T-Cost, we used a continuous optimization method to determine the best-possible reward that could be applied to each block of actions by applying a common translation in action space to all actions (Fig. 2D, top panel). The T-Cost was quantified as the difference between this optimized cost and the original cost. For the N-Cost, we used continuous optimization to determine the best-possible reward obtainable by scaling the distance of each action from the mean across all actions. The N-Cost was quantified as the difference between this optimized cost and the original cost. For the C-Cost, we shuffled the pairings between different selected release velocities and angles to find the pairing that yielded the best-possible reward, which we identified through a greedy pair-swapping approach. The C-Cost was quantified as the difference between this optimized cost and the original cost.

### De Novo Learning Model

The model for the de novo learning task follows very similarly to the model for the skittles task. In this task, however, we aim to learn a policy that varies based on the target location (represented by the state *s*). We assume that targets appear at a fixed distance *d* but at a variable angle *s* from the start location. Participants must learn appropriate displacements of the left and right hands *a*_*L*_ and a_*R*_ in order to bring the cursor to the target, so that the action *a* in each trial is a 2-d vector: *a* = (a_*L*_, *a*_*R*_). We assumed that the final location of the cursor was given by *c* = (a_*L*_, *a*_*R*_). c = (a_L_,*a*_*R*_). The reward function was set to be equal to the negative of the absolute distance between the cursor and target location, i.e.

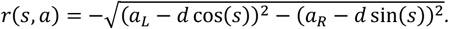

The movement policy specifies a distribution of displacements for the left and right hands that is now dependent on target angle *s*. We again assume that this distribution is Gaussian and that the m ean displacements of the left and right hands are represented as a sum of Gaussian basis functions, i.e.

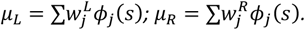

Since the target could appear in any direction, we used a circular basis function taking the shape of the von Mises distribution, taking the form

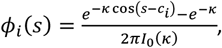

where κ is a concentration parameter (analogous to the reciprocal of the variance of a Gaussian) and ***I***_8_ is the modified Bessel function of the first kind of order zero. We included 31 basis functions with centers *c*_*i*_ equally spaced around the circle, and set κ = 5, corresponding to a basis function width of approximately 25^°^. In this case, the policy mean is parametrized by *w*^ti^ and *w*^?^. The policy gradient update for these parameters is given by:

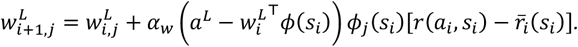

For the covariance, we assume a single parameter, corresponding to the log-variance, that applies equally to left and right hand actions, and is independent of target direction, i.e.

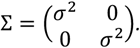

Finally, since performance could be drastically different depending on the state, we maintained a state-dependent average reward (i.e. a value function) for each possible target state *s*. The update rule for σ is then given by

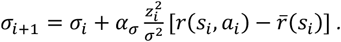

Policy-gradient RL models initialized with randomized weights had an initial directional much higher than human learners and initial learning was quite slow. To better mimic human performance, we initialized the model by matching the initial policy to human participants’ early performance (blocks 2-3 of 35). In human experiments, participants were able to make online corrections and each trial only ended when the cursor reached the target. We couldn’t base the initialized policy on participants’ endpoint data. Instead, we constructed projected endpoints based on initial velocities of the left and right hands (taken 100ms after movement onset). We rescaled these initial velocities to convert them into projected endpoints, scaling all trials by a fixed amount (per participant) determined such that the mean cursor displacement matched the distance to the target (12cm). We used least-squares regression with L_2_ regularization (ridge regression, *α* = 1) to determine initial policy weights. The residual variance around this learned policy was used to initialize *v*_*i*_ for each action dimension. The policy parameters were normalized based on the mean initial standard deviation for the two task-relevant action dimensions. Lastly, we used a separate, slower learning rate for the variance parameters, *α*_FG_ = 0.02 to better align with human performance, which showed slower improved compared to when the policy-gradient variance learning rate was matched to that of the mean (*α* = 0.1).

### Arc task model

In the arc task, participants make rapid movements to guide a cursor through a narrow arc from a start location to an end location^16^. A feature of behavior in such tasks is the presence of highly regular sub-movements occurring with a highly regular period of 130 ms. Following a model proposed by Guigon^21^, we model this behavior using an optimal feedback control model with intermittently-updated movement goals. We model action selection in this task as selecting an appropriate sequence of abstract movement goals to achieve task success.

We model movements with duration 780 ms, so as to consist of 6 sub-movements of 130ms duration each. During each sub-movement, we assumed that movements pursued a fixed spatial goal **g**^*i*^ following an optimal controller with a horizon of 280 ms (a value which has been shown to account for the sub-movement structure of human movements). We modeled the wrist as a dynamical system with state **x** comprising position *x*, velocity *x*?, and two muscle states *m*_1_ and *m*_2_:

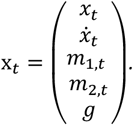

Here, *g* is the current goal location, which is also incorporated into the state, which allowing the endpoint cost to be expressed as *J* = **x**^6^*Q* **x**, with

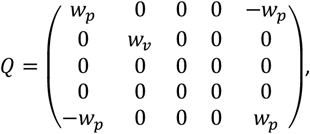

where *w*_H_ and *w*_9_ are cost weights penalizing the squared deviation from the target, and squared velocity at the endpoint of the movement. The control cost is given by *J*_**u**_ = **u**^6^**u**.

The dynamics are given by

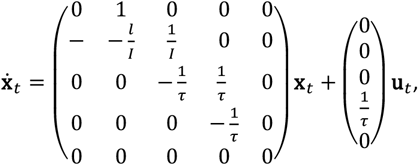

expressed more compactly as 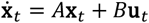 the above equation, ***I*** is the inertia of the wrist, *l* is viscosity, and τ is a time constant related to muscle activation.

We extend this 1-d model to two dimensions by doubling the number of states, and changing the matrices to be block-diagonal, i.e.

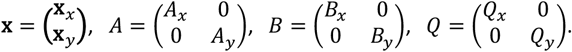

We discretized this problem using matrix exponentials, using a discretization timestep of 0.001s, and solved for the optimal time-varying control gains *L*_7_ using the RicaN equations (see Todorov), so that the motor commands on timestep *i* were given by **u**_*i*_ = -*L*_*i*_**x**_*i*_ **u**_i_.

To simulate a movement for a given sequence of *N* goals

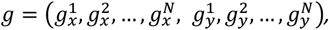

we set the goal states in according to the initial goal location, and simulated the first 130ms of a 280ms movement using the optimal control gains *L*_*i*_. After each 130ms, the goal states were updated and a new movement was simulated starting from the end state of the prior movement, and resetting the control gains to *L*_1_for the new submovement.

For the policy gradient optimization, we took participants’ action in each trial to be the specifaction of the sequence of goals **g** driving the movement. We assume that participants’ policy was given by a multivariate Gaussian distribution:

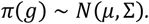

We initialized µ as a sequence of equally-spaced points along the center of the arc. We initialized Σ as a diagonal matrix with equal variance for all dimensions, i.e. Σ = σ_8_***I***.

We assumed that the reward function for successful movements comprised a number of components. First, we penalized deviations from the center of the arc:

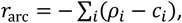

Where *ρ*_*i*_ reflects the radial position of the cursor on timstep *i*. We also included a term to encourage smooth trajectories by penalizing movement jerk:

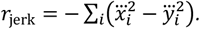

We also included a state reward at the end of the movement equivalent to the negative of the costs used in the optimal control problem 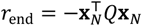he total reward was then given by:

Policy updates were as per equations 4 and 5.

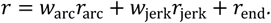

All simulations were developed in python. Code for generating all simulations is available on the author’s GitHub page: github.com/adrianhaith/PolicyGradientSkillLearning.

## Acknowledgements

Thanks to Eric Grießbach, John Krakauer and Joshua Cashaback for helpful comments on the paper.

